# EZH2 Inhibition Reshapes 3D Chromatin Architecture to Induce Immunogenic Phenotype in Small Cell Lung Cancer

**DOI:** 10.64898/2026.01.26.701784

**Authors:** Sana Parveen, Ramesh Adhinaveni, Kun Fang, Lavanya Choppavarapu, Meijun Du, Gustavo Leone, Navonil de Sarkar, Victor Jin, Hui-Zi Chen

## Abstract

**Background:** The histone methyltransferase EZH2, enzymatic core of the trimeric polycomb repressive complex 2 (PRC2), has been shown to promote small cell lung cancer (SCLC) survival through epigenetic silencing of multiple targets including Class I MHC molecules (*HLA-A/B*) and DNA repair factors (*SLFN11*). Treatment of SCLC cells with EZH2 inhibitors *in vitro* can reactivate expression of these genes and result in therapeutic response to immune checkpoint inhibition (ICI) and chemotherapy. Here, we investigate the impact of EZH1/2 dual inhibition on 3D chromatin structure and its relationship to transcriptional regulation in neuroendocrine (NE) SCLC.

**Results:** Employing Micro-C, a micrococcal nuclease-based 3D genome mapping technique, we show that EZH1/2 inhibition with Valemetostat induced significant changes at multiple genome organizational levels (compartment, topological associated domain, and chromatin loop) without incurring cell death in NE SCLC. Alterations in 3D genome permissive for transcriptional activation were correlated with increased chromatin accessibility (ATAC-sequencing) and expression of target genes (transcriptome profiling). Known transcription factor motif discovery revealed enrichment of non-NE motifs (e.g., *REST*) in regions with gained chromatin accessibility in Valemetostat-treated cells, consistent with results from gene set enrichment analysis demonstrating NE to non-neuroendocrine lineage shift. Notably, EZH1/2 inhibition reactivated Class I MHC expression by facilitating enhancer-promoter looping.

**Conclusion:** Our results demonstrate that repression of a subset of EZH2 targets including Class I MHC genes is affected through modulation of 3D genome structure to the level of chromatin looping and further support clinical investigation of EZH2 inhibition in boosting therapeutic efficacy of ICI in SCLC patients.

## BACKGROUND

Small cell lung cancer (SCLC) is a rare aggressive bronchogenic lung cancer (a reported incidence of 4.7 cases per 100,000 individuals in 2021) that constitutes ∼15% of new lung cancer cases diagnosed globally each year[1]. Most patients present with metastatic or extensive stage SCLC (ES-SCLC), which has an estimated 5-year overall survival (OS) rate of ∼12%[1]. Although SCLC is initially very treatment sensitive, almost all patients will relapse and demonstrate resistance to subsequent therapies. Currently, none of the FDA approved treatments for ES-SCLC requires biomarker testing, including the newly approved DLL3/CD3 bi-specific T cell engager (BiTE) therapy[2]. SCLC is now recognized as a molecularly heterogeneous cancer that encompasses neuroendocrine (NE) and non-neuroendocrine (NNE) subtypes arising from distinct progenitor cells[3, 4], with subtype-specific therapeutic vulnerabilities[5]. Furthermore, preclinical studies have demonstrated epigenetic-mediated subtype plasticity or phenotype conversion in SCLC as a major mechanism of therapeutic resistance[6, 7].

In contrast to oncogene-driven non-small cell lung cancer (NSCLC), the most prevalent genomic alterations in smoking-induced SCLC are inactivating mutations in tumor suppressor genes including *TP53*, *RB1* and *PTEN*[8–10]. Multiple mouse models have demonstrated the central role of concurrent *Tp53* and *Rb1* loss in SCLC tumorigenesis[11]. Additionally, earlier whole genome analyses of primary SCLC tumors and cell free DNA from patients showed recurrent mutations in chromatin-remodeling genes including *CREBBP*/*EP300* and *KMT2D*[8, 10, 12], implicating crucial roles for chromatin organization and its transcriptional impact on SCLC initiation or progression. Indeed, *Crebbp* loss in mice accelerated tumorigenesis in the classic *Trp53/Rb1*-deficient SCLC model and sensitized tumors to inhibition with histone deacetylases *in vivo*[13]. Finally, a unique subset of SCLC with intact *RB1* and *TP53* has been identified in never or light smokers, whose tumors were shown to develop from recurrent chromothripsis leading to amplification of cell cycle regulators[14].

The mammalian Polycomb Repressive Complex 2 (PRC2) is composed of three key proteins EED, SUZ12, and EZH2 and is a master regulator of differentiation by precisely coordinating the timely expression of developmental genes[15]. EZH2 is a histone methyltransferase (HMT) that catalyzes the tri-methylation of histone 3 lysine 27 (H3K27me3), which is a chromatin mark associated with compacted chromatin and transcriptional silencing[15]. Earlier research indicated high *EZH2* expression and therefore a potential dependency on its function in human SCLC [16]. Subsequent studies in SCLC patient-derived xenografts (PDXs) and cell lines have elucidated that EZH2 silences numerous genes necessary for therapeutic response to both chemotherapy (*SLFN11*) and immunotherapy (*TAP1, HLA-A/B/C*)[7, 17]. Pharmacologic inhibition of EZH2 restored antigen presenting capacity in SCLC cells, sensitizing them to anti-PD-1 blockade[17]. The same immunogenic phenotype was also seen when SCLC cells were treated with an inhibitor of lysine-specific demethylase 1 (LSD1)[18, 19]. Finally, genetic ablation of *Kdm6a*, a H3K27 histone demethylase, in a murine SCLC model altered chromatin accessibility and promoted SCLC subtype switching[6]. Altogether, the current data supports epigenome deregulation as one of the principal mechanisms that drives SCLC subtype plasticity and therapeutic resistance.

In this original report, we conducted multi-omic analyses of an NE SCLC cell line (NCI-H146) to further investigate the mechanisms of how EZH2 governs oncogenic transcription in SCLC. Along with whole transcriptome and chromatin accessibility profiling, we performed Micro-C sequencing[20, 21], a micrococcal nuclease (MNase)-based three-dimensional (3D) genome mapping technique with superior spatial resolution derived from the original high-throughput chromatin conformation capture (Hi-C) assay, to reveal that the expression EZH2 target genes (e.g., *HLA-B/C*) necessary for therapeutic response in SCLC is regulated through long-range chromatin looping that normally enables enhancer-promoter contact. Our results showcase a precise mechanism of how EZH2 exerts its oncogenic function in SCLC.

## RESULTS

### EZH2 is a Main Epigenetic Transcriptional Silencer in NE SCLC

We performed an exploratory analysis of *EZH2* expression in 22 types of solid cancer cell lines from the CCLE in cBioPortal and determined that SCLC (n=50) demonstrated the highest median *EZH2* expression (**Fig. 1A**). We next evaluated the expression of all PRC2 components (*EZH2, SUZ12,* and *EED*) in the same fifty SCLC cell lines, which showed high expression of *EZH2* and *SUZ12* regardless of subtype classification (**Fig. 1B**)[22]. Based on the association with specific accessory proteins, mammalian PRC2 can be classified into two independent subcomplexes, PRC2.1 and PRC2.2, which share overlapping target genes in mouse embryonic stem cells[15, 23]. Interestingly, we detected higher expression of PRC2.1 complex proteins (*PHF1* and *PHF19* encoding individual Polycomb-like proteins) than PRC2.2 complex proteins (*AEBP2* and *JARID2*) in SCLC cell lines (**Fig. 1B**). We further explored the expression of a previously published list of genes (n=61) enriched for the repressive histone mark H3K27me3 and whose promoters were bound by SUZ12 as determined by chromatin immunoprecipitation in a SCLC cell line Lu130[24]. We found decreased expression of most of these genes in the CCLE SCLC cell lines (**Fig. S1**), consistent elevated PRC2 function in SCLC.

**Figure 1.**
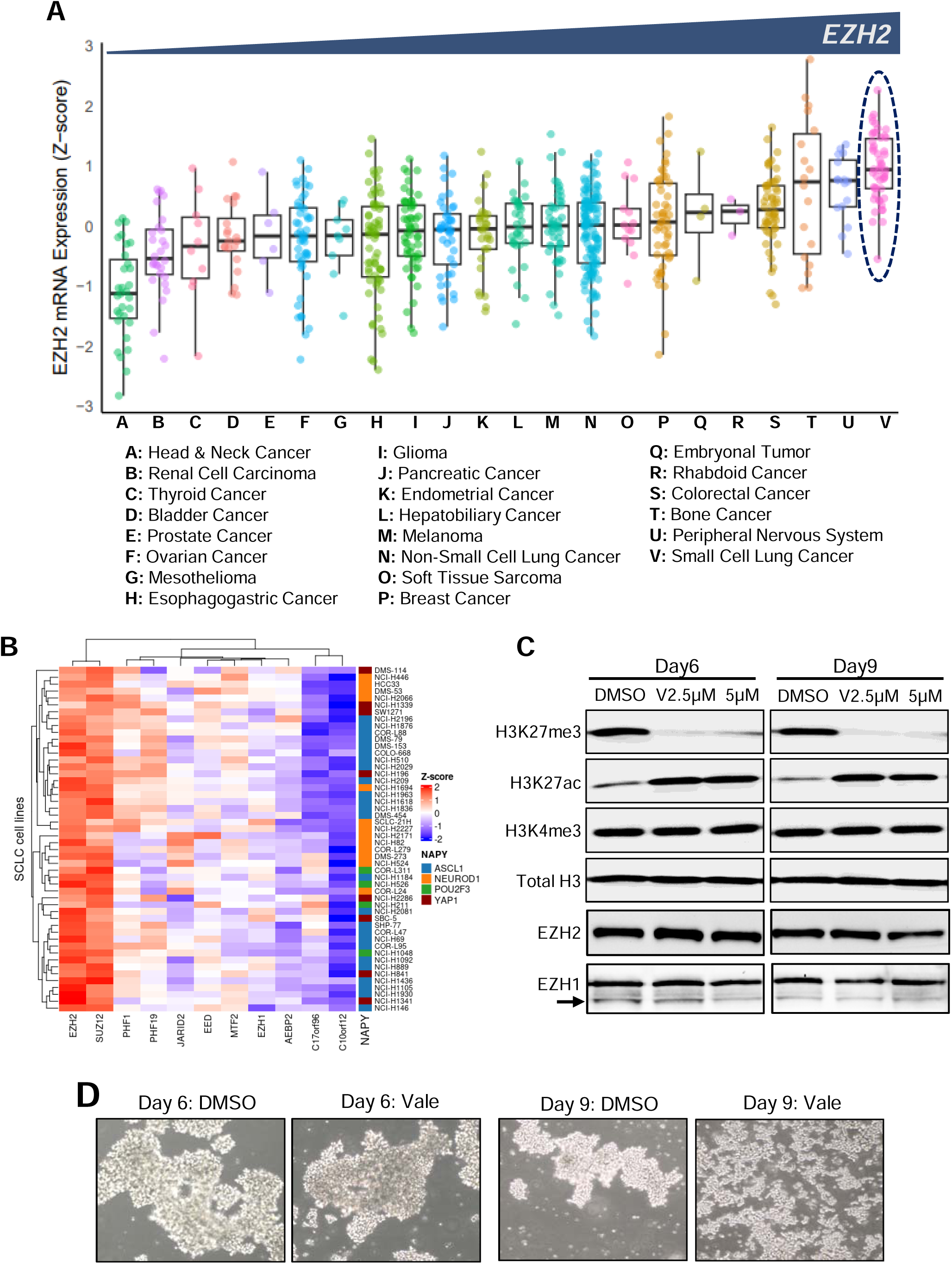
EZH2 is effectively inhibited by Valemetostat in small cell lung cancer (SCLC). **(A)** EZH2 mRNA expression in 22 solid cancer types from the CCLE database, ordered by increasing median expression (left to right). SCLC cell lines (n=50) exhibited the highest median *EZH2* expression. **(B)** RNA-seq heatmap demonstrating expression of various PRC2 components in CCLE SCLC cell lines (n = 50). Cell lines are annotated according to their molecular subtype (ASCL1, NEUROD1, POU2F3, and YAP1)[22]. **(C)** Immunoblot analysis of H146 cells treated with Valemetostat (2.5µM and 5µM) or DMSO for 6 and 9 days. Reduction of H3K27me3 levels confirms the enzymatic inhibition of EZH1/2 by Valemetostat. Active enhancer-specific H3K27 acetylation (H3K27ac) increased, while active promoter-specific H3K4 trimethylation (H3K4me3) showed no change. Total EZH2 and EZH1 protein levels remained consistent throughout day 9 of experiment. Arrow points to band corresponding to EZH1 protein. Total H3 levels were probed as control. **(D)** Representative bright-field microscopy images of H146 cells demonstrating change in growth pattern (from clusters in suspension to floating/semi-adherent monolayer) after treatment with 2.5µM Valemetostat. Images were captured at 10x magnification.

After confirming RNA expression of PRC2 components in SCLC cell lines, we performed immunoblot to evaluate the effects on protein stability of direct EZH2 pharmacologic inhibition in a well-characterized NE SCLC cell line (H146) that lacks Class I MHC expression due to PRC2-mediated epigenetic silencing[17, 25]. While EZH2-specific inhibitors are available (*e.g.*, GSK126), we specifically chose Valemetostat, a dual EZH1/EZH2 inhibitor, to account for potential functional compensation by EZH1 when EZH2 activity is lost[26]. Valemetostat currently has regulatory approval for treatment of T cell malignancies in Japan and is in clinical trial testing for various solid tumors globally. We treated H146 cells with increasing concentrations of Valemetostat, which led to a significant decrease in the repressive chromatin mark H3K27me3 that is associated with silenced enhancers by day 6 of treatment compared with DMSO control (**Fig. 1C**). By day 9, the reduction of H3K27me3 was even more profound (**Fig. 1C**). As global H3K27me3 levels are dramatically reduced, the mark of active enhancer regions as represented by H3K27 acetylation (H3K27ac) was simultaneously increased[26] (**Fig. 1C**). The mark of transcriptionally active promoters, H3K4me3, was not affected between control and treated cells. Finally, we observed stable protein levels of both EZH2 and EZH1 in H146 cells treated with Valemetostat, supporting the fact that the widespread changes in enhancer histone modifications were due to PRC2 functional inhibition rather than decreased protein levels (**Fig. 1C**). Finally, continuous Valemetostat exposure induced a change in the growth pattern of H146 cells, from clusters in suspension to sheet-like monolayers with partially adherent properties (**Fig. 1D**), suggesting subtype switching since non-neuroendocrine SCLC subtypes have been observed to preferentially grow as adherent cells[27]. This altered growth pattern was consistently reproducible across multiple independent experiments.

### EZH1/2 Inhibition Induces Inflammatory Response and Non-Neuroendocrine (NNE) Differentiation in SCLC

We performed transcriptome sequencing to interrogate the effects on global gene transcription of EZH1/2 dual inhibition in H146 cells. At day 9 post-treatment with Valemetostat and DMSO control, H146 cells were collected for total RNA extraction followed by Illumina sequencing. As shown via heatmap and volcano plot, approximately 75% (3,352/4,453) of significantly differentially expressed genes (DEGs) were upregulated with dual EZH1/2 inhibition (**Fig. 2A-B**), consistent with the known function of PRC2 in mediating gene silencing in the basal state. We selected several upregulated genes and validated their increased expression by quantitative RT-PCR (**Fig. 2C**). The expression of *REST*, which suppresses NE differentiation[28], and *SLFN11*, which was previously shown to be a direct EZH2 target[7], were significantly upregulated by RT-PCR. The expression of *CD44*, which is a marker of epithelial to mesenchymal differentiation[27], was also significantly increased. We next applied Gene Set Enrichment Analysis (GSEA) to our DEGs and showed a positive enrichment for genes in Antigen Processing and Inflammatory Response pathways (**Fig. 2D**), consistent with results from prior studies[17]. Furthermore, we noted a positive enrichment of Non-Neuroendocrine gene signature and decreased enrichment of Neuroendocrine signature, supporting NE to NNE transition induced by functional inhibition of PRC2 (**Fig. 2D**). Notably, a substantial proportion of SUZ12-bound genes (37/61) also demonstrated significantly increased expression upon Valemetostat exposure in H146 cells (**Fig. S2**). Collectively, these findings indicate that PRC2 has two important functions in SCLC: to epigenetically silence genes involved in innate anti-tumor immune response and to maintain NE differentiation via suppression of genes such as *REST*.

**Figure 2.**
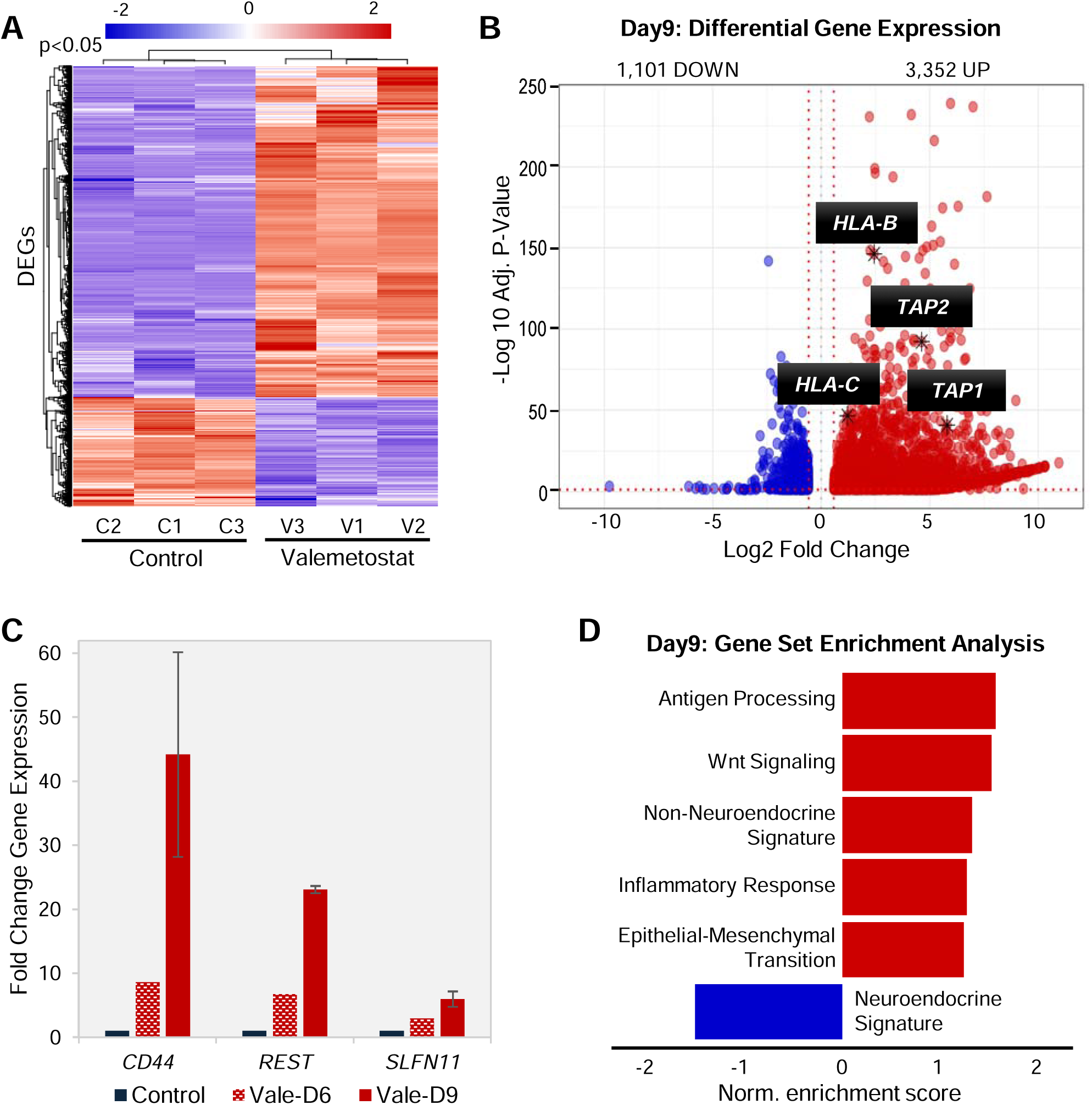
Transcriptome sequencing reveals that EZH1/2 inhibition induces inflammatory response and non-neuroendocrine (NNE) differentiation in SCLC. **(A)** Heatmap showing differentially expressed genes (DEGs) in H146 cells following treatment with Valemetostat for 9 days relative to vehicle control. Independent triplicate samples were included per timepoint and condition. **(B)** Volcano plot illustrating DEGs between Valemetostat day 9 and control H146 cells, with a cutoff of Log_2_Fold Change≥∣0.58∣ and adjusted p-value <0.05. Antigen processing/presentation genes are highlighted (*e.g.*, *TAP1*) and show significantly induced expression with EZH1/2 inhibition. **(C)** Quantitative RT-PCR analysis of selected genes at day 6 and 9, including the positive control gene *SLFN11*, the cancer stem cell marker *CD44*, and the non-neuroendocrine marker *REST*. *SLFN11* is included as a known target gene repressed by EZH2 in SCLC [7]. **(D)** Gene Set Enrichment Analysis (GSEA) of DEGs between Valemetostat day 9 and control H146 cells, demonstrating positive enrichment of Antigen Processing and NNE signatures and negative enrichment of NE signature.

### EZH1/2 Inhibition Increased Chromatin Accessibility Globally Correlating with Transcriptional Activation

To determine the chromatin landscape underlying the transcriptional changes induced by pharmacologic PRC2 inhibition, we performed ATAC-sequencing of Valemetostat and DMSO-treated H146 cells at day 9 and conducted differential accessibility analysis using CoBRA workflow[29]. Heatmap of differentially accessible genomic regions (**Fig. 3A**) revealed that Valemetostat treatment led to increased chromatin accessibility of ∼32,000 genomic regions (gained, **Fig. 3B**) and decreased chromatin accessibility of ∼9,900 genomic regions (lost, **Fig. 3B**). The 9,900 genomic regions with lost accessibility in Valemetostat-treated cells are the same as regions with increased accessibility in DMSO-treated cells. We then mapped the differentially accessible regions to genomic locations corresponding to the following features: gene promoter, intron/exon, distal intergenic regions, and 5’/3’-untranslated regions (UTR) (**Fig. 3C**). This analysis showed that EZH1/2 inhibition resulted in approximately 10% gain in accessible ‘distal intergenic’ regions, correlating with a loss in accessible ‘intronic’ genomic regions, compared with DMSO control (**Fig. 3C**). As enhancer elements are often located in distal intergenic regions, these results suggest that long-range interactions, such as promoter-enhancer interactions facilitated by chromatin looping, are responsible for the transcriptional changes seen with PRC2 inhibition.

**Figure 3.**
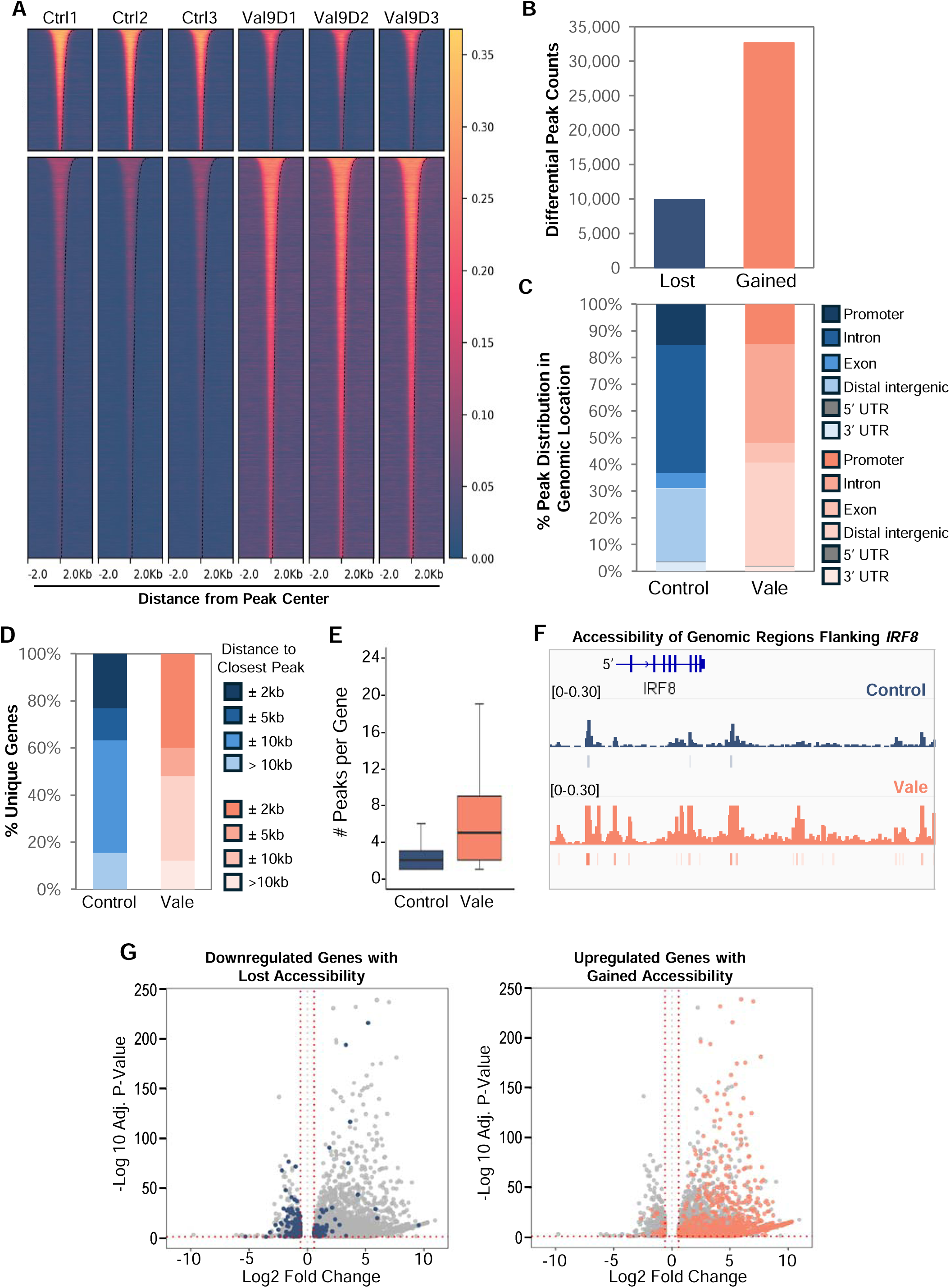
EZH1/2 Inhibition increased chromatin accessibility globally as shown by ATAC-sequencing, correlating with transcriptional activation. **(A)** Peak-centered heatmap (±2kb) displaying differential accessible regions following day 9 Valemetostat treatment in H146 cells compared to control, with Log_2_Fold Change≥∣1∣ and adjusted p-value <0.05. Independent triplicate samples were included per timepoint, and condition and differential accessible regions were performed using CoBRA workflow. **(B)** Bar plot showing unique differential peak counts (Log_2_Fold Change≥∣1∣ and adjusted p-value <0.05) in day 9 Valemetostat treated and control H146 cells. Lost: unique peaks detected in control cells. Gained: unique peaks detected in treated cells. **(C)** A stacked bar plot illustrating the distribution of differential peaks across various genomic regions in day 9 Valemetostat treated and control H146 cells. Genomic regions include promoters, introns, exons, 5’ and 3’-untranslated regions (UTRs), and distal intergenic regions. **(D)** Proximity of differential peaks to Transcription Start Sites (TSSs) using a distance-based peak binning of ATAC-seq peaks in day 9 Valemetostat treated and control H146 cells. For each gene, only the single closest accessible peak to its respective TSS is to classify TSS-peak distance as indicated in the legend. **(E)** Box plot depicting median number of peaks per gene in which closest peaks were detected ± 2kb from TSS in day 9 Valemetostat and control samples. Center line is median, while box shows IQR. **(F)** Integrative Genomics Viewer (IGV) example of *IRF8* locus, a representative example, demonstrating gained chromatin accessibility (increased number of ATAC-seq peaks in day 9 Valemetostat-treated cells compared to control) in regions proximal and distal to the gene. This gain in chromatin accessibility corresponded to increased gene expression. **(G)** Volcano plot showing the correlation between changes in chromatin accessibility (±2 kb from TSS) and gene expression. Dark blue dots represent downregulated genes with decreased accessibility in day 9 Valemetostat-treated cells; orange dots represent upregulated genes with increased accessibility in day 9 Valemetostat-treated cells.

We next performed combined analysis of transcriptome and ATAC-seq data from H146 cells. We curated a list of genes by assigning a single closest ATAC-seq peak to a nearest gene and segregated this list into categories based on peak distance from Transcriptional Start Site (TSS): ±2 kilobases (kb), ±5 kb, ±10 kb, and >10kb (**Fig. 3D**). We found that a higher percentage (∼40%) of genes in EZH1/2-inhibited H146 cells exhibited increased accessibility ±2 kb of TSS (**Fig. 3D**); this is almost twice the percentage (∼20%) of genes belonging to the same classification than in control cells. Furthermore, of the genes with ATAC-seq peak assignment ±2 kb of TSS, we then examined whether more peaks were called in genomic regions surrounding and throughout these genes. We found that Valemetostat-treated H146 cells showed overall higher number of peaks (median: 5 peaks/gene) surrounding or distributed throughout a gene than DMSO control (median: 2 peaks/gene) (**Fig. 3E**). As an example, Figure 3F shows IGV view of *IRF8*, which demonstrated gained accessibility ±2 kb of TSS and further upstream in distal intergenic regions (**Fig. 3F**).

Finally, we overlapped DEGs with genes having ATAC-seq peaks ±2 kb of TSS. Volcano plot visualization shows that most downregulated DEGs correlated with decreased accessibility (dark blue dots) while most upregulated DEGs correlated with increased accessibility (light orange dots) (**Fig. 3G**). Tools like HOMER have been developed to identify potential transcription factor (TF) binding sites within differentially accessible chromatin regions[30]. Using HOMER, we conducted known TF motif discovery using as input peaks that were gained and lost in H146 cells following treatment with Valemetostat (**Fig. 3B**). Notably, we identified a significant enrichment for REST binding motifs in peaks gained upon Valemetostat treatment (-log_10_ p-value = 19). This finding, combined with the observed transcriptional upregulation of *REST* (**Fig. 2C**), shows that PRC2 inhibition promotes NNE differentiation of SCLC cells. On the other hand, CTCF motifs were highly enriched in control-specific peaks (lost peaks upon Valemetostat treatment) (**Fig. S3A**), consistent with CTCF’s known role in acting as an insulator that dampens enhancer-promoter interactions[31]. Altogether, these results indicate that pharmacologic PRC2 inhibition leads to a dynamically open chromatin state enabling transcriptional reprogramming in NE SCLC.

### EZH1/2 Inhibition Leads to Reorganization of Higher Order 3D Chromatin Structure (Compartments and Topological Associating Domains) in SCLC

Micro-C is a 3D genomic sequencing technology, also known as chromatin conformation capture assay, that can map chromosome interactions with resolution down to the nucleosome level[21]. Micro-C can be applied to chromatin structure analysis, structural variation detection, and analysis of enhancer-promoter interactions[20]. The 3D genome is organized into compartments (of megabase-scale chromatin), topological associating domains (TADs), chromatin loops, and at the smallest scale nucleosome-nucleosome interactions[32]. We performed Micro-C in H146 cells after 9 days of Valemetostat or DMSO treatment and subsequent compartment level analysis was conducted using ‘dchic’ at 100 kb resolution[33]. In our data analysis, we found that EZH1/2 inhibition resulted in a transition of many B compartments (transcriptionally inactive) to A compartments (transcriptionally active) as demonstrated in the Sankey plot (**Fig. 4A**). The A/B compartments can be further divided into sub-compartments (*e.g.*, A0, A1, and A2)[34], and we showed that A compartments remained as A compartments or transitioned from A0 to even more transcriptionally active A1 or A2 sub-compartments with EZH1/2 inhibition (**Fig. 4A** and **S4A**).

**Figure 4.**
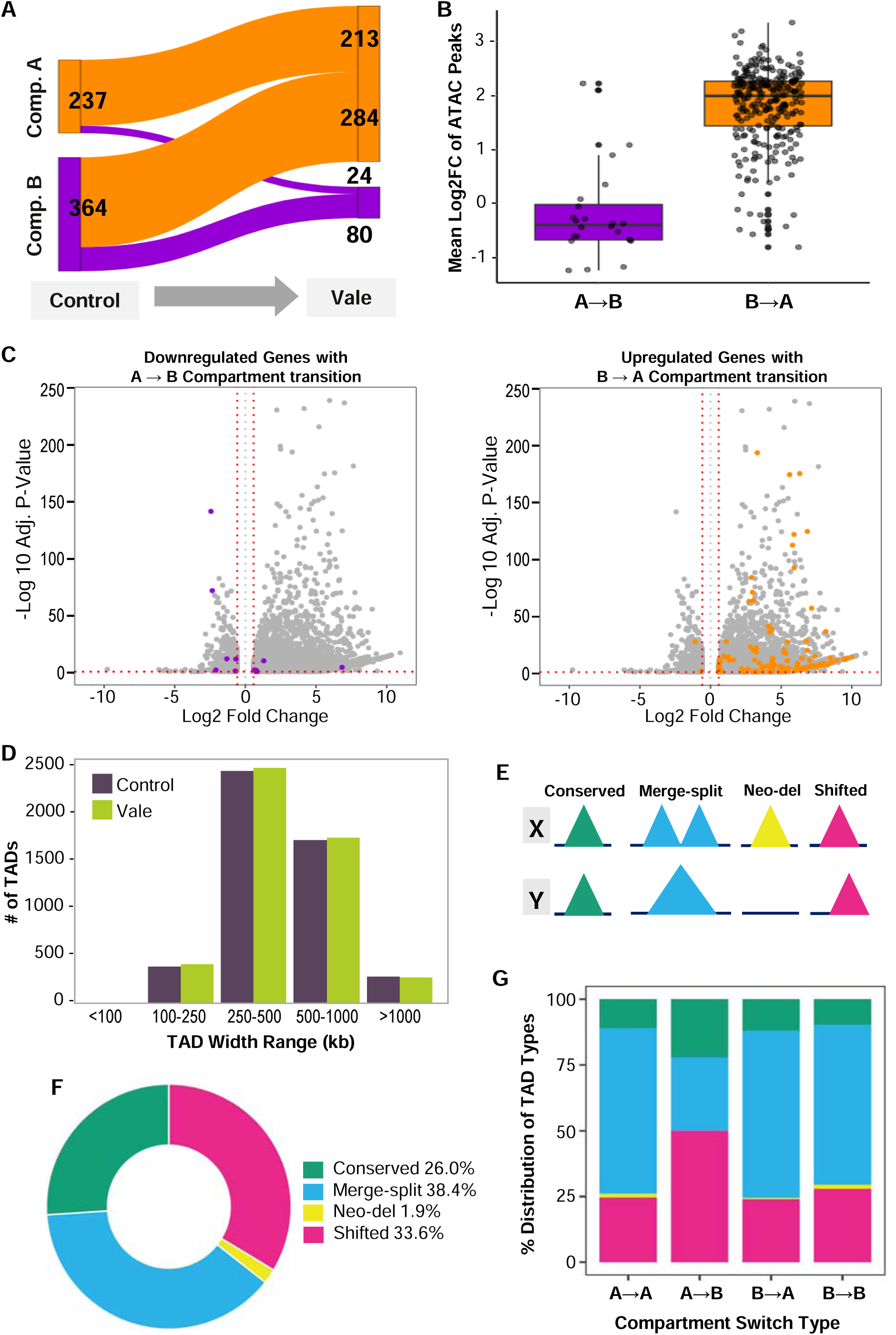
EZH1/2 Inhibition leads to reorganization of higher order 3D chromatin structure in SCLC. Changes in A/B compartment and topologically associating domain (TAD) landscapes following 9 days of treatment with 2.5 µM Valemetostat, compared to a vehicle treated controls, were analyzed using ‘dcHiC’ and ‘cooltools’. **(A)** Sankey plot illustrating A/B compartment transition of 601 differential compartments between day 9 Valemetostat-treated and control samples. Compartment A is depicted as an orange node, and Compartment B as a purple node. The connecting arc represents the switching (A→B or B→A) or matching (A→A or B→B) compartment transition. **(B)** Box plot showing the mean differential accessibility scores of differentially accessible regions (detected by ATAC-seq) that overlap with compartment flipping regions (A→B and B→A) in day 9 Valemetostat-treated cells. B→A transitions demonstrate significantly higher mean differentially accessibility score. Center line indicates the median, and box represents IQR. **(C)** Volcano plots highlighting DEGs located within the A→B and B→A transitioning compartments. Most upregulated genes correlated with B→A (closed to open) compartment switch. **(D)** Grouped bar plot showing the distribution of TADs with indicated widths (in kb) identified in control and Valemetostat-treated samples using ‘cooltools’. The mean TAD width in both groups was ∼500 kb. **(E)** Representative illustration of the different types of TAD modification that may be detected between two theoretical samples X and Y. **(F)** Donut plot showing frequency of the four categories of TAD modification in combined TADs from day 9 Valemetostat-treated and control samples. The least common modification was ‘neo-del’ while the most common was ‘merge-split’. **(G)** Stacked bar chart representing overlap between the four types of TADs and differential A/B compartment transitions. Color scheme is consistent with Figure 3E.

We next reasoned that ATAC-seq accessibility could serve as a proxy for assessing chromatin compaction in regions undergoing differential compartment transitions. Therefore, we extracted genomic coordinates for A→B and B→A compartment transitions induced by EZH1/2 inhibition and overlapped these with ATAC-seq peaks. Our results showed that B→A transitions, as well as A→A sub-compartment transitions, exhibited significantly higher ATAC-seq peak accessibility scores compared to A→B or B→B transitions (**Fig. 4B** and **S4B**), supporting increased chromatin accessibility in the A compartments. The B→A shift also correlated with increased transcriptional activity, as DEGs in the genomic regions corresponding to B→A transitioning demonstrated overall increased expression (**Fig. 4C**).

To investigate how PRC2 inhibition impacts TAD integrity in H146 cells, we employed insulation score-based TAD calling using ‘cooltools’, with 10kb bin size and 250kb sliding window[35]. TADs exceeding 1.5 Mb were excluded, yielding 4,792 TADs in control samples and 4,856 in Valemetostat-treated samples. The overall distribution of TAD width was similar between the two groups, with a mean TAD width of ∼500 kb (**Fig. 4D** and **S5A**), consistent with prior reported TAD widths in human sample[36]. TADs were classified into four categories, conserved, shifted, merge-split, and neo-del (**Fig. 4E**), using established methods[37, 38]. This yields 26% conserved, 38.4% merge-split, 33.6% shifted, and 1.9% neo-del in Valemetostat-treated vs. control cells (**Fig. 4F**). As shifted TAD was the second most abundant category suggesting minor boundary repositioning while largely preserving domain architecture, we next investigated the potential functional consequences of these shifts on the adjacent TADs. Boundary regions, separating adjacent TADs, were extracted (**Fig. S5B**), and differential boundary analysis was conducted using the ‘quaich’ workflow. Differential boundaries were identified based on insulation score ratios between conditions and/or boundary gains arising from shifted TAD endpoints, neo-del TAD termini, or TAD splitting. Of the 4,886 boundaries in control samples, 44% (2,148/4,886) were differential (gained or stronger insulation relative to Valemetostat-treated sample); similarly, 45% (2,202/4,939) of boundaries in treated samples were differential (gained or stronger relative to control). Paired TAD analysis at differential boundary regions revealed frequent proximity of shifted TADs to merge-split and neo-del regions (**Fig. S5C**). Collectively, these findings indicate that while ‘shifted’ TADs largely preserve domain architecture via boundary repositioning and modest insulation alterations, they impact neighboring domains, leading to modified intra- and inter-TAD interactions and context-dependent chromatin reorganization upon EZH1/2 inhibition.

As TADs are the basic folding units that are organized into higher-order A/B compartments[39], we next evaluated whether the distribution of TAD categories (**Fig. 4E**) varied across the differential A/B compartments in Valemetostat-treated cells relative to control. We extracted genomic loci mapping to differential A/B compartments and plotted the distribution of TAD reorganization induced by EZH1/2 inhibition within these regions (**Fig. 4G**). From this analysis, we found that ‘Merge-split’ and ‘Shifted’ TADs occurred with highest frequencies across all compartment switch types observed (blue and magenta, **Fig. 4G**), suggesting that TAD boundary rearrangement was the most prevalent reorganization event. Collectively, these findings demonstrate that Valemetostat-induced epigenetic reprogramming modulate higher order chromatin structure to influence gene expression in SCLC.

### EZH1/2 Inhibition Enables Enhancer-Promoter Contact of Antigen Presentation Genes via Chromatin Looping in SCLC

Chromatin loops are formed in cis between two spatially separated genomic regions on the same chromosome, with each loop having two ‘anchor regions’ (**Fig. S6A**), and facilitate the physical juxtaposition of important genetic elements to regulate gene expression[40]. Using an established algorithm MUSTACHE[41], we performed loop calling with 5 kb resolution setting and identified approximately 16,000 chromatin loops configured in cis in both control and Valemetostat-treated H146 cells. The mean size of loops called was ∼250 kb (**Fig. 5A**), consistent with what has been previously reported in human cells[40]. To focus our analysis on biologically relevant loops while accounting for the discrete nature of 5 kb binning during loop calling, we applied a slightly adjusted size cutoff of 255 kb (250 kb + 5 kb) and excluded loops exceeding this threshold, retaining 8,471 loops for the remaining analyses in this section. Of the 8,471 chromatin loops, 4,087 (48.2%) were conserved or detected in both control and Valemetostat-treated cells (**Fig. S6B**), while 1,793 (21.2%) and 2,591 (30.6%) chromatin loops were identified as unique in control and Valemetostat-treated cells, respectively (**Fig. 5B** and **S6B**).

**Figure 5.**
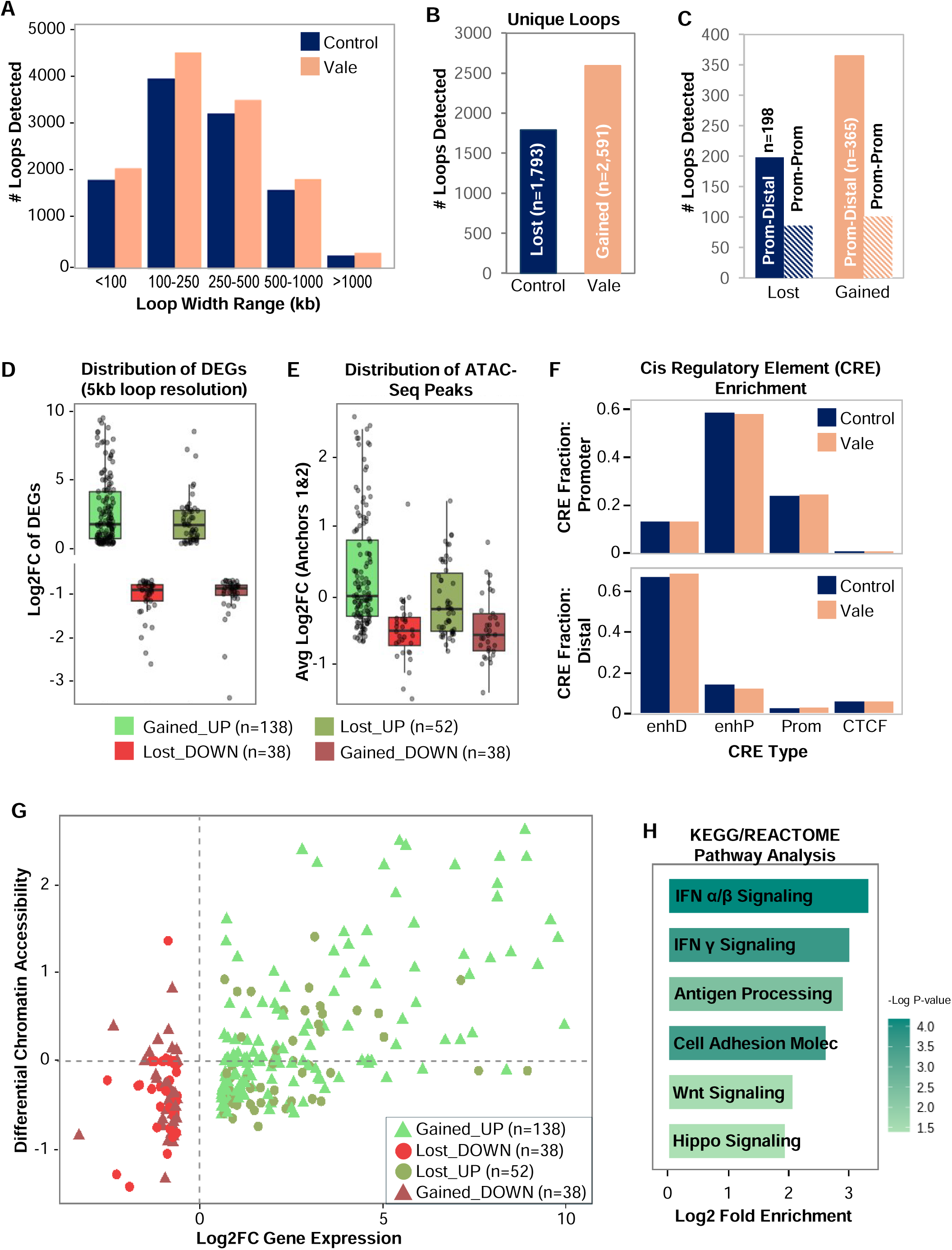
EZH1/2 inhibition facilitate chromatin looping involving promoters and putative enhancer regions to activate gene expression. **(A)** Grouped bar plot illustrating the distribution of chromatin loop widths in day 9 Valemetostat-treated and control samples. **(B)** Bar plot displaying number of unique chromatin loops ≤255 kb identified in day 9 Valemetostat-treated and control samples. These unique loops are classified as ‘lost’ (present only in control) or ‘gained’ (present only in Valemetostat-treated samples). **(C)** Bar plot showing the counts of unique loops (lost or gained) sub-classified by genomic location of anchors: Promoter-Promoter (P-P) or Promoter-Distal (P-D). **(D)** Box-and-whisker plot showing expression (upregulation or downregulation) of DEGs in gained and lost loops in day 9 Valemetostat-treated cells. The highest number of upregulated DEGs (138) was detected in gained loops (‘Gained_UP’). Center line indicates the median, and box represents IQR. Number of genes in each category is indicated in parentheses. **(E)** Box-and-whisker plot depicting the distribution of differential accessibility scores (ATAC-seq peaks) at the two anchors of gained and loops in the same four categories as defined in Figure 4D. **(F)** Bar plots illustrating the enrichment of Cis Regulatory Elements (CREs) in promoter (top plot) and distal (bottom plot) anchors in unique promoter-distal loops in control and Valemetostat-treated cells. 5 kb anchors were intersected with ATAC-seq peaks to identify accessible sub-regions. Human CREs in the ENCODE registry were then overlapped with these anchor-restricted accessible peaks. enhD, enhancer-distal; enhP, enhancer-proximal; Prom, promoter. **(G)** Scatter plot integrating RNA-seq, ATAC-seq, and Micro-C data. X-axis represents gene expression, while Y-axis represents differential chromatin accessibility. ▴ indicate genes where a gained loop (P-P or P-D) was detected; ● indicate genes where a lost loop (P-P or P-D) was detected. Categories of loop status (gained or lost) and expression of DEGs are as defined in Figure 4D and 4E. **(H)** Pathway enrichment analysis (KEGG and REACTOME) using DAVID[57], performed on DEGs associated with gained chromatin loops.

We began by annotating and mapping the genomic locations (i.e. promoter, gene body, and distal elements) of both anchor regions of all unique chromatin loops (**Fig. S6C**). Each anchor region of a chromatin loop represents a genomic region spanning 5 kb. We defined ‘promoter’ region as 5 kb upstream or 2 kb downstream of a gene TSS, while ‘distal element’ (signifying potential enhancer regions) was defined as 250 kb upstream or downstream of TSS. With these definitions, our loop anchor annotations were consistent with previously published report, in which ∼15-20% of chromatin loops were formed between ‘promoter-promoter’ and ‘promoter-distal’ anchor points (gray and darker purple bars, **Fig. S6C**)[34], representing loops with high potential to affect transcriptional regulation. In Valemetostat-treated cells, we detected a higher absolute number of loops with ‘promoter-distal’ anchor regions (365 loops) compared with control cells (198 loops), suggesting increased enhancer-promoter interactions as the basis for transcriptional changes seen with EZH1/2 inhibition (solid bars, **Fig. 5C**).

To assess whether chromatin looping induced by EZH1/2 inhibition affects gene expression in H146 cells, we performed integrative analysis of Micro-C and transcriptome data. We selected chromatin loops with ‘promoter-promoter’ and ‘promoter-distal’ anchor regions and evaluated their overlap with DEGs from RNA-seq (**Fig. 5D**). Consistent with expectations, newly formed or ‘gained loops’ in Valemetostat-treated cells were associated with increased gene expression (green boxplot, **Fig. 5D)** while ‘lost loops’, or those detected in DMSO but not Valemetostat-treated cells, were associated with decreased gene expression (red boxplot, **Fig. 5D**). Interestingly, a subset of genes exhibited the opposite trend, where ‘lost loops’ correlated with increased gene expression and ‘gained loops’ with decreased expression (dimmed green/red boxplots, **Fig. 5D**). This latter unexpected association between looping and gene expression pattern suggests that certain distal genetic elements may function as transcriptional silencers rather than enhancers, and their disrupted interaction with promoters may lead to gene activation rather than repression[42].

To investigate the influence of chromatin accessibility on looping-induced differential gene expression in H146 cells, we calculated differential ATAC-seq accessibility scores within anchor regions of ‘gained’ and ‘lost loops’ (focusing again on ‘promoter-promoter’ and ‘promoter-distal’ loops) in Valemetostat-treated cells (**Fig. S6D**). We detected the highest accessibility scores in ‘gained loops’ associated with up-regulated genes (green boxplot, **Fig. 5E**). ‘Lost loops’ associated with down-regulated genes demonstrated lower accessibility scores (red boxplot, **Fig. 5E**). Finally, we also examined the distribution of accessibility scores in anchor regions of loops involving up or down-regulated genes in control cells (dimmed green/red boxplots, **Fig. 5E**). Overall, we conclude from this analysis that looping-mediated transcriptional activation of genes, whether in Valemetostat-treated or control cells, is accompanied by increased chromatin accessibility.

To further investigate whether the distal anchor in ‘promoter-distal’ loops may function as a putative enhancer element, we performed an analysis whereby all accessible (as determined through ATAC-seq) anchor regions of unique ‘promoter-distal’ loops in Valemetostat-treated (365) and control H146 cells (198) were mapped against a defined set of Cis-Regulatory Elements (CREs) from the ENCODE registry[43]. These CREs included the following categories: enhancer-proximal, enhancer-distal, promoter, and CTCF. In both Valemetostat-treated and control cells, high proportions (>60%) of ‘distal’ anchor regions of chromatin loops mapped to ‘enhancer-distal’ elements as defined by ENCODE (enhD, **Fig. 5F**). Approximately 10% of ‘distal’ anchor regions also mapped to ‘enhancer-proximal’ elements (enhP, **Fig. 5F**). Altogether, ∼70% of ‘distal’ anchor regions mapped to putative enhancer elements in the genome. On the other hand, the largest fraction of ‘promoter’ anchor regions mapped to ENCODE-defined ‘enhancer-proximal’ regions (∼50%) followed by ‘promoter’ regions (∼20%) (enhP and Prom, **Fig. 5F**).

In summary, our results demonstrate that EZH1/2 inhibition in H146 cells enabled the formation of chromatin loops between putative proximal as well as distal enhancer elements and gene promoters, which further demonstrate a correlation with enhanced chromatin accessibility and transcriptional activation (**Fig. 5G**). Using DEGs associated with ‘gained loops’ in Valemetostat-treated cells as input, KEGG/REACTOME analysis detected significant enrichment of Interferon (IFN) Signaling, Antigen Processing, as well as Wnt and Hippo signaling pathways (**Fig. 5H**). Perturbed Wnt signaling was previously shown to drive chemotherapy resistance in SCLC cell lines[44], while the Hippo axis was associated with certain metastatic behaviors in SCLC *in vitro*[45]. IGV visualizations of gained or lost loops in control and Valemetostat-treated cells are presented in Figure 6. Unique ‘promoter-distal’ loops involving *DNAJC10* and *SMC2* were detected in control H146 cells (**Fig. 6A**), while unique ‘promoter-distal’ loops involving *TACSTD2* and *FZD8* corresponding with their increased gene expression (Log_2_FC 1.53 and 2.49, respectively) were detected in Valemetostat-treated cells (**Fig. 6B**). *TACSTD2* encodes a cell-surface glycoprotein (TROP2) that is currently being investigated as a therapeutic target in *de novo* SCLC patients[46]. Additionally, ‘promoter-promoter’ and ‘promoter-distal’ loops involving *HLA-B/C* were detected in Valemetostat-treated cells only (**Fig. 6B**), which showed a time-dependent increase in *HLA-B* and *HLA-C* expression through differential gene expression analysis (**Fig. 2B**) and RT-PCR (**Fig. 6C**). Finally, cell-surface expression of Class I MHC induced by EZH1/2 inhibition was confirmed using flow cytometry with a pan-HLA antibody and appropriate isotype control. H146 cells were treated with 5 µM of EZH2-specific inhibitors (GSK126 and Tazemetostat) or 2.5 µM Valemetostat; by day 9, pan-HLA surface expression was highest in cells treated with Valemetostat (blue, **Fig. 6D**).

**Figure 6.**
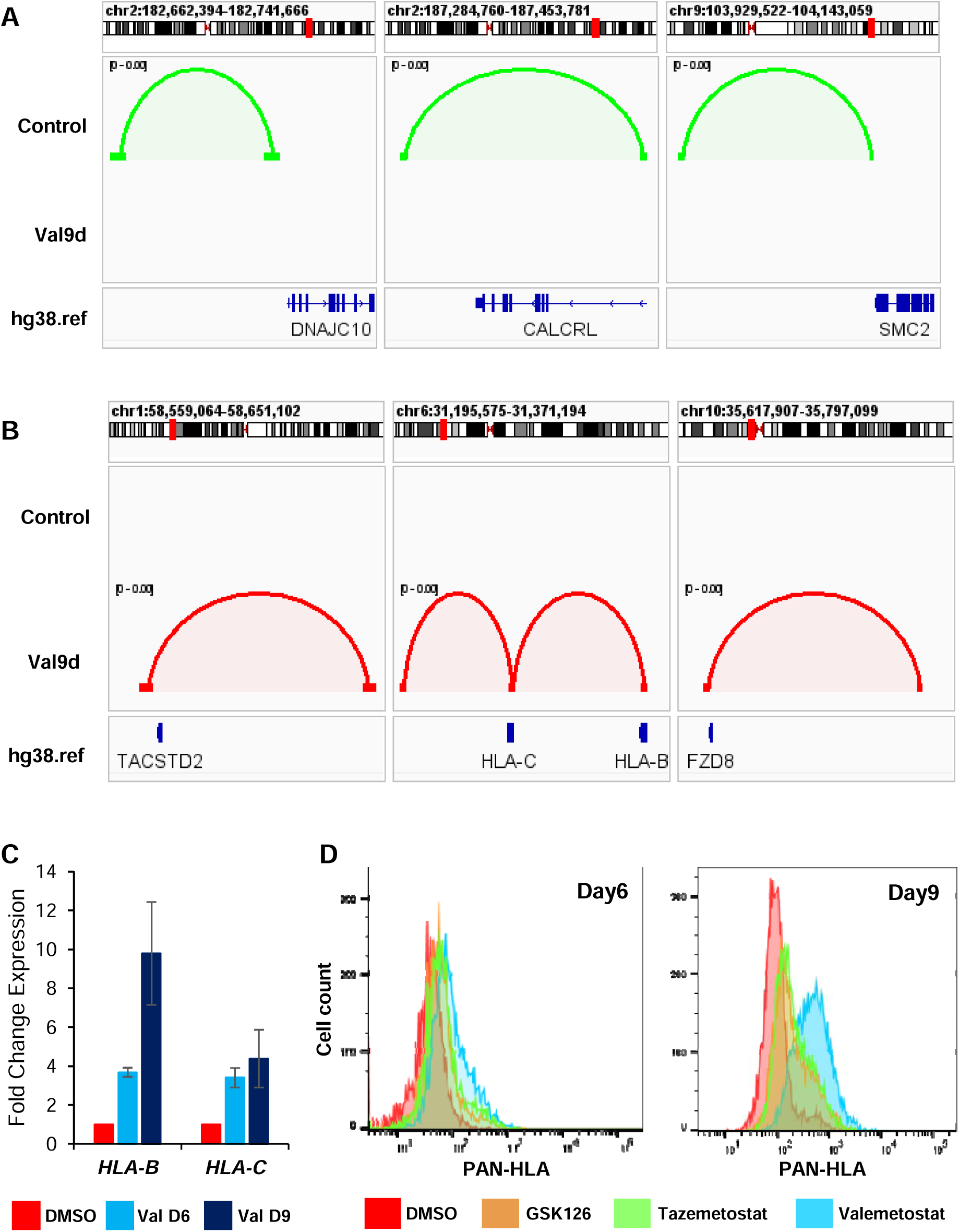
EZH1/2 inhibition enables long-range interaction via looping to activate target gene transcription in SCLC. IGV representation of **(A)** lost and **(B)** gained loops identified in day 9 Valemetostat-treated cells. Loops are classified as ‘lost’ (green, present only in control) or ‘gained’ (red, present only in Valemetostat-treated samples). *TACSTD2*, *HLA-B/C*, and *FZD8* were amongst DEGs demonstrating increased expression with gained P-P or P-D loops in day 9 Valemetostat-treated cells. Chromosomal location of each gene is highlighted in red; location of each gene as shown within reference human genome hg38. **(C)** Quantitative RT-PCR of *HLA-B* and *HLA-C* expression in day 6 and day 9 Valemetostat-treated and control samples. **(D)** Flow cytometry analysis of surface Class I MHC expression as detected by pan-HLA antibody in H146 cells treated with control, EZH2-specific (5µM GSK126 and Tazemetostat) and EZH1/2 dual (2.5µM Valemetostat) inhibitors at indicated time points.

## DISCUSSION

Disrupted integrity of the 3D genome at multiple hierarchical levels (compartment, TAD, chromatin loop) is a core underlying mechanism of human cancer development[47]. The original 3D chromatin conformation capture technique Hi-C has been employed in tissue or cell lines to investigate acquired therapeutic resistance and tumor heterogeneity in breast cancer, acute myeloid leukemia, and colorectal cancer[48]. Recently, Choudhuri *et al.* performed Hi-C in serial PDX models – (1) treatment-naïve and (2) post-treatment with Olaparib/Temozolomide – from a single SCLC patient as part of a broader investigation to discern the role of *MYC* in driving cross-resistance of relapsed SCLC[49]. Overall, however, there is sparse knowledge regarding state of the 3D genome in SCLC and its evolution under therapeutic pressure. To interrogate 3D genomic alterations in NE SCLC induced by pharmacologic PRC2 inhibition, we performed Micro-C using MNase, which cuts genomic DNA between two nucleosomes to create nucleosome-sized fragments for proximity ligation, thereby achieving higher spatial resolution than Hi-C[20, 21]. We combined Micro-C with chromatin accessibility (ATAC-seq) and transcriptome profiling in NE SCLC to thoroughly unveil the impact of complete PRC2 inhibition, which was achieved using a dual EZH1/2 inhibitor Valemetostat. Our results demonstrate the importance of PRC2-maintained chromatin structure in silencing genes critical for NE differentiation and immunogenicity in SCLC.

EZH2 has long been identified as a key oncogenic protein with classical function as an HMT in SCLC[16, 24]. In SCLC patients who received treatment with either single (anti-PD-1) or doublet (anti-PD-1 and anti-CTLA4) immunotherapy, high *EZH2* expression in tumors correlated with worse overall survival[50]. Previously, Mahadevan *et al.* showed that EZH2-specific inhibition with GSK126 rescued expression of antigen processing/presentation genes (APGs) including *TAP1/2* and *HLA-A/B/C*[17]. Furthermore, functional studies including pan-HLA I immunopeptidome analysis demonstrated gained ability of GSK126-treated SCLC cells to process and present endogenous peptides[17]. Indeed, this important study propelled our current multi-omic investigation, which now provides an additional dimension to the mechanism of how EZH2 coordinates repression of target genes including APGs based on multi-level modulation of chromatin structure. At the first level of A/B compartmentalization, EZH1/2 inhibition produced preferential compartmental shifts from closed (B) to open (A) formation. Valemetostat-induced A compartments were associated with an overall gain in chromatin accessibility, leading to increased expression of gene subsets. At the next level of TADs, EZH1/2 inhibition significantly altered the TAD landscape, leading to an abundance of TADs (with a mean size of ∼500 kb) defined by shifted boundaries and formation of new TADs through merge/split of pre-existing TADs. CREs (*e.g.*, enhancers and promoters) within a TAD are more likely to interact with one another than with elements outside the TAD due to presence of insulating TAD boundaries enriched for CTCF and cohesin proteins[51]. Our conclusion from these analyses is that EZH1/2 inhibition led to considerable higher order chromatin structure alterations to account for the differential expression of a proportion of PRC2-regulated genes in SCLC.

Chromatin looping is a form of long-range interaction that brings together distal CREs such as enhancers into proximity of target gene promoters, leading to more robust transcriptional activation (though the opposite has also been observed due to existence of ‘silencer’ elements[42]). Inactivating mutations in histone-modifying enzymes such as *KMT2D* and *CREBBP*/*EP300* are recurrently identified in human SCLC tumors and cell lines [8, 10, 12], supporting a critical role for sustained enhancer integrity in suppressing SCLC development. *KMT2D* encodes a member of the lysine-specific methyltransferase family (MLL) of proteins that mediate mono-methylation of H3K4 (H3K4me1), an epigenetic histone mark specifically associated with active enhancers. Heterozygous or homozygous KMT2D mutations in SCLC cell lines lead to reduced H3K4me1 levels and deregulated gene expression[52]. Loss-of-function mutations in the histone acetyl transferase CREBBP and its paralog EP300 in SCLC likely result in globally reduced H3K27 acetylation (H3K27ac), another mark associated with active enhancers. These observations altogether support disruption of looping-mediated enhancer-promoter interactions as a key mechanism to repress transcriptional activation of known and novel genes with anti-tumor functions in SCLC.

In our chromatin looping analysis (loop size restricted to < 255 kb), we detected a gain in chromatin loops with ‘promoter-distal’ and ‘promoter-promoter’ anchor regions when EZH1/2 was inhibited. The anchor regions of gained loops mapped to ENCODE-defined CREs including proximal/distal enhancer regions, which were further characterized by increased chromatin accessibility. Furthermore, when all three levels of data (transcriptome, chromatin accessibility, and 3D chromatin structure) are combined, we detect a positive correlation amongst gained loops, increased accessibility, and transcriptional activation (**Fig. 5G**). The classical MHC-I molecules (HLA-B/C) were among the most notable EZH2 target genes whose expression was determined to be regulated by looping in our multi-omic dataset. Tumor-specific downregulation of MHC-I occurs in numerous human cancers via mechanisms such as direct gene mutations, antigen depletion, and transcriptional modulation[53]. Based on our Micro-C data, we further refine a model whereby the primary mechanism of MHC-I downregulation in SCLC is through transcriptional modulation due to restricted physical contact between the more proximally located cis-acting promoter elements (*i.e.*, enhancer A and interferon-stimulated response element (ISRE)[54]) and distal enhancer elements. This physical contact is prohibited in SCLC cells by a functional PRC2, innately disabling antigen presentation and subsequent anti-tumor immune response. With PRC2 inhibition, we show that the expression of IFN genes is induced – which can activate classical MHC-I expression via proximal promoter elements like ISRE[54] – and looping further occurs to enable distal enhancer-promoter contact, resulting in HLA-B/C RNA and protein expression in our model. Indeed, our pathway analysis revealed that the expression of IFN genes themselves is also regulated by looping.

## CONCLUSION

In summary, we performed a novel 3D genome mapping technique Micro-C to investigate how EZH2 orchestrates oncogenic transcription in SCLC through modulation of chromatin structure. Our results show that EZH1/2 dual inhibition triggered transcriptional reprogramming in SCLC that led to an effect on differentiation (NE to NNE shift), increased IFN signaling, and increased expression of APGs (and other clinically relevant targets such as *TACSTD2*) due to looping-mediated enhancer-promoter contact. These results further support efforts to refine therapeutic PRC2 inhibition, in rational combination with immune checkpoint blockade, to elicit durable responses in ES-SCLC patients.

## METHODS

### Cell lines

The human small cell lung cancer cell line H146 was obtained from Dr. Anish Thomas’ group and cultured in RPMI-1640 medium supplemented with 10% fetal bovine serum (FBS) and 1X Penicillin-Streptomycin (Penstrep). Cell lines were authenticated by short tandem repeat (STR) profiling and routinely tested for mycoplasma contamination using the InvivoGen MycoStrip Detection Kit.

### Pharmacological treatment

H146 cells were treated with Valemetostat (MedChemExpress, HY-109108A), Tazemetostat (Selleckchem, S7128), GSK126 (Abcam, ab269816) or DMSO control for 9 days with media being replenish every third day. Cells were harvested on day 6 or day 9, washed twice with ice-cold PBS and the resulting cell pellets were stored in −80°C.

### Immunoblotting

Cell pellets were thawed on ice and lysed in blue juice lysis buffer (250 mM Tris, pH 6.8, 20% glycerol, 8% SDS) containing 2-mercaptoethanol. Equal protein amounts from the lysates were separated by denaturing SDS-PAGE (12% or 8% gels) and then transferred to PVDF membranes. Membranes were blocked for 1 hour with 5% non-fat milk in Tris-Buffered Saline with 0.1% Tween 20 (TBS-T) and subsequently incubated overnight at 4°C with primary antibodies diluted in either 5% BSA or 5% non-fat milk in 1X TBST. The following day, membranes were washed three times with 1X TBST and incubated with appropriate HRP-conjugated secondary antibodies. Protein detection was performed after a final three washes with 1X TBST using the Clarity Western ECL Substrate (Bio-Rad, 1705061). All primary and secondary antibodies were purchased from Abcam, unless otherwise specified. The H3K27ac antibody was obtained from Diagenode (C15410196). Other antibodies used in this study were as follows: Anti-EZH1 (ab289887), Anti-EZH2 (ab186006), Anti-Histone H3 (tri methyl K27) (ab6002), Anti-Histone H3 (ab1791), Anti-Histone H3 (tri methyl K4) (ab8580), Goat Anti-Rabbit IgG H&L (HRP) (ab6721), and Rabbit Anti-Mouse IgG H&L (HRP) (ab6728).

### RNA isolation and quantitative Realtime PCR

Total RNA was extracted using the RNeasy Kit (Qiagen, 74104) following the manufacturer’s instructions. One microgram of RNA was reverse transcribed into cDNA using the High-Capacity cDNA Reverse Transcription Kit (Applied Biosystems™, 4368814). Quantitative PCR (qPCR) for *GAPDH, HLA-B, HLA-C, SLFN11, CD44, and REST* was performed using Power SYBR™ Green PCR Master Mix (Applied Biosystems™, 4368702). *GAPDH* served as the internal control, and relative gene expression was quantified using the ΔΔCT method. The primer sequence used for RT-PCR were *CD44* Forward- 5’ CGC AGC CTG GGG ACT CTG 3’, *CD44* Reverse- 5’ CGA GAG ATG CTG TAG CGA CCA 3’; *GAPDH* Forward- 5’ATG GGG AAG GTG AAG GTC G 3’, *GAPDH* Reverse- 5’ GGG GTC ATT GAT GGC AAC AAT A 3’; *HLA-B* Forward- 5’ GAT GGC GAG GAC CAA ACT CA 3’, *HLA-B* Reverse- 5’ CTC CGA TGA CCA CAA CTG CT 3’; *HLA-C* Forward- 5’ CCT GAG CTG GGA GCC ATC 3’, *HLA-C* Reverse- 5’ CAG CTA GGA CAA CCA GGA CA 3’; *REST* Forward- 5’ AAC TCA TAC AGG AGA ACG CCC 3’, *REST* Reverse- 5’ TAG AGG CCA CAT AAC TGC ACT G 3’, and *SLFN11* Forward- 5’ GGC CCA GAC CAA GCC TTA AT 3’, *SLFN11* Reverse 5’ CAC TGA AAG CCA GGG CAA AC 3’.

### Flow cytometry for surface HLA expressions

H146 cells were cultured in the presence of the indicated inhibitors (5 µM GSK126, 5 µM Tazemetostat, or 2.5 µM Valemetostat) alongside a vehicle control for up to 9 days, with drug-containing media replenished every third day. Flow cytometry samples were collected on day 6 and day 9, and staining was performed on the same day. For each sample, 1 million cells were washed twice with 1X PBS and subjected to live/dead staining using Zombie NIR™ (BioLegend, 423105) followed by staining with FITC-conjugated PAN-HLA (BioLegend, 311404) or isotype control antibodies (BioLegend, 400208). Following staining, cells were fixed using Cyto-Fast™ Fix/Perm Buffer (BioLegend, 426803) and analyzed for surface PAN-HLA expression on BD LSRFortessa™ cell analyzer. Data were analyzed using FlowJo software where FITC signal was captured exclusively from live cells using appropriate gating strategies.

### RNA-seq library preparation and analysis

The whole-transcriptome sequencing library was prepared using Illumina TruSeq stranded mRNA library prep kit according to the Manufacturer’s protocol and subjected to 100bp paired-end sequencing on NovaSeq 2000 platform.

All RNA sequencing data was analyzed using nfcore/rnaseq pipeline version 3.14.0[55]. Briefly, the workflow began with quality control using FastQC, followed by adapter trimming with Trim Galore. Reads were then aligned to the hg38 genome using STAR with the GENCODE v43 annotation GTF file. Gene quantification was performed with Salmon and differential gene expression analysis with DESeq2.

### ATAC-seq library preparation and analysis

ATAC seq for control and Valemetostat treated H146 samples were performed according to manufacturer’s recommendations (Active motif cat # 53150). Briefly, nuclei were isolated from 50k viable cells using ATAC-seq lysis buffer followed by tagmentation reaction. Later, the next-gen sequencing-ready library was purified, QC, and sent for 100bp paired-end sequencing on the NovaSeq 6000 platform (Illumina). ATAC-seq analysis was performed using nfcore/atacseq pipeline (version 2.1.2) (https://nf-co.re/atacseq/2.1.2)[55]. Briefly, paired-end reads were aligned to the human genome hg38 using Bowtie2. The blacklist and PCR duplicates were removed from the aligned reads and the chromatin accessible peaks were called using MACS2 in narrowPeak mode with default parameter. To perform the differential peak accessibility, coverage of narrow peak between control and valemetostat treated samples were extracted using multiBamSummary of deepTools (https://deeptools.readthedocs.io/en/latest) followed by DESeq2 analysis as described in CoBRA workflow[29].

### Micro-C library preparation

Micro-C profiling was performed using Dovetail’s Micro-C kit following the manufacturer’s instructions. Briefly, 1 million cells were frozen in −80°C and subsequently crosslinked at room temperature using a combination of disuccinimidyl glutarate (DSG) and formaldehyde. The crosslinked samples underwent micrococcal nuclease (MNase) digestion to achieve optimally fragmented chromatin. Quality control (QC) of chromatin digestion was assessed using a fragment analyzer on a small sample volume, targeting a mononucleosome population of 40–70%. Once QC was confirmed, the remaining digested chromatin was subjected to proximity ligation on chromatin capture beads, followed by adapter ligation. To selectively amplify long-range ligation events, the ligated fragments were captured on streptavidin beads and subjected to PCR amplification using HotStart PCR ready mix with unique sample-specific indexing. The amplified products were then size-selected using SPRIselect beads. Final next-generation sequencing (NGS) libraries were prepared, purified, and sequenced using paired-end 150 bp reads on an Illumina NovaSeq 6000 platform.

### Micro-C Data processing and analysis

Paired-end FASTQ files were aligned to the hg38 reference genome using the BWA-MEM algorithm with the parameters −5SP -T0. Valid ligation events were extracted using pairtools (https://pairtools.readthedocs.io/en/latest/) with the parameters --min-mapq 40 --walk-policy 5unique --max-inter-align-gap 30, and the resulting pairsam files were processed to remove PCR duplicates. The final filtered .pairs files were generated using the pairtools split function. The processed pairs files were used to generate multi-resolution contact matrices with cooler (https://cooler.readthedocs.io/en/latest). For downstream analysis, dcHiC was employed for differential A/B compartment analysis at 100kb resolution[33]. Cooltools insulation was used to identify differentially enriched topologically associating domains (TADs) with a 10 kb bin size and a 250 kb window[35]. TADs were classified into conserved, shifted, merge-split, and neo-del categories based on genomic overlap between Control and Valemetostat detected TADs, following established methods[37, 38]. Conserved TADs exhibited 100% reciprocal overlap; shifted TADs showed one-to-one correspondence with boundary repositioning of ≤5 bins and ≥80% reciprocal overlap; merge-split TADs encompassed one-to-many (split) or many-to-one (merge) relationships with ≥80% coverage of overlapping segments; and neo-del TADs lacked detectable overlap between conditions. Differential boundary analysis was performed using quaich workflow (https://github.com/open2c/quaich). Chromatin loops were detected using Mustache at 5 kb resolution[41]. Both common (shared) and condition-specific chromatin loops were classified using the approach described by Dozmorov *et al*[56].

## ABBREVIATIONS

APGs: Antigen processing/presentation genes
ATAC: Assay for Transposase Accessible Chromatin
CCLE: Cancer Cell Line Encyclopedia
CREs: Cis regulatory elements
DEG: Differentially expressed gene
ENCODE: Encyclopedia of DNA Elements
HLA: Human Leukocyte Antigen
MHC: Major Histocompatibility Complex
MNase: Micrococcal nuclease
NE: Neuroendocrine
NNE: Non-neuroendocrine
PRC2: Polycomb Repressor Complex 2
SCLC: Small cell lung cancer
TAD: Topological Associated Domain

## ETHICS DECLARATIONS

Not applicable

## DATA AVAILABILITY

EZH2 expression data from solid cell lines were downloaded from the cBioPortal database (https://www.cbioportal.org/). The RNA expression data of small cell lung cancer cell lines were downloaded from Cancer Cell Line Encyclopedia (CCLE) website (https://sites.broadinstitute.org/ccle/datasets). The sequencing data generated in this study are deposited in the NCBI Gene Expression Omnibus under the following accession numbers. RNA-seq, ATAC-seq are available under accession number GSEXXXX. Micro-C data is available under accession number GSEYYYY.

## CODE AVAILABILITY

The scripts used for all data analyses and figure generation in this study are available in a public GitHub repository at https://github.com/sanamcw/EZH2i_microC_analysis under an MIT license.

## AUTHOR CONTRIBUTIONS

SP, RA, and HZC designed and performed experiments. SP performed bioinformatics analyses; KF, LC, MD, and VJ assisted in bioinformatic analyses. SP, VJ and HZC analyzed data. SP and HZC wrote the manuscript. GL, NS, and VJ revised and edited the manuscript.

## ACKNOWLEDGEMENTS

We thank Neshatul Haque and Raul Urrutia for their valuable discussions throughout this study. We thank Angela Mathison, Jaime Wendt Andrae, and Michael Tschannen for their contributions to RNA and ATAC sequencing, which was performed at the Mellowes Center for Genomic Sciences and Precision Medicine at the Medical College of Wisconsin. We also thank Zhao Lai, from the Genome Sequencing Facility (GSF) core at Greehey Children’s Cancer Research Institute (Greehey CCRI) at UT Health San Antonio (UTHSA), for sequencing the micro-C data. We are grateful to Anish Thomas and his group for providing the H146 cell line. We also thank Galina Petrova for her expert guidance and training at the Flow Cytometry Core, Children’s Research Institute, Medical College of Wisconsin. Additionally, we appreciate the technical support provided by Myriam El Khawand, Mia Truong, and the Cantata Bio team in optimizing Micro-C nuclease digestion and interpreting quality-control metrics. This work was supported in part by computational resources and technical assistance from the Research Computing Center at the Medical College of Wisconsin.

