## Supplementary material for "EZH2 Inhibition Reshapes 3D Chromatin Architecture to Induce Immunogenic Phenotype in Small Cell Lung Cancer": N/A

### Slide 1
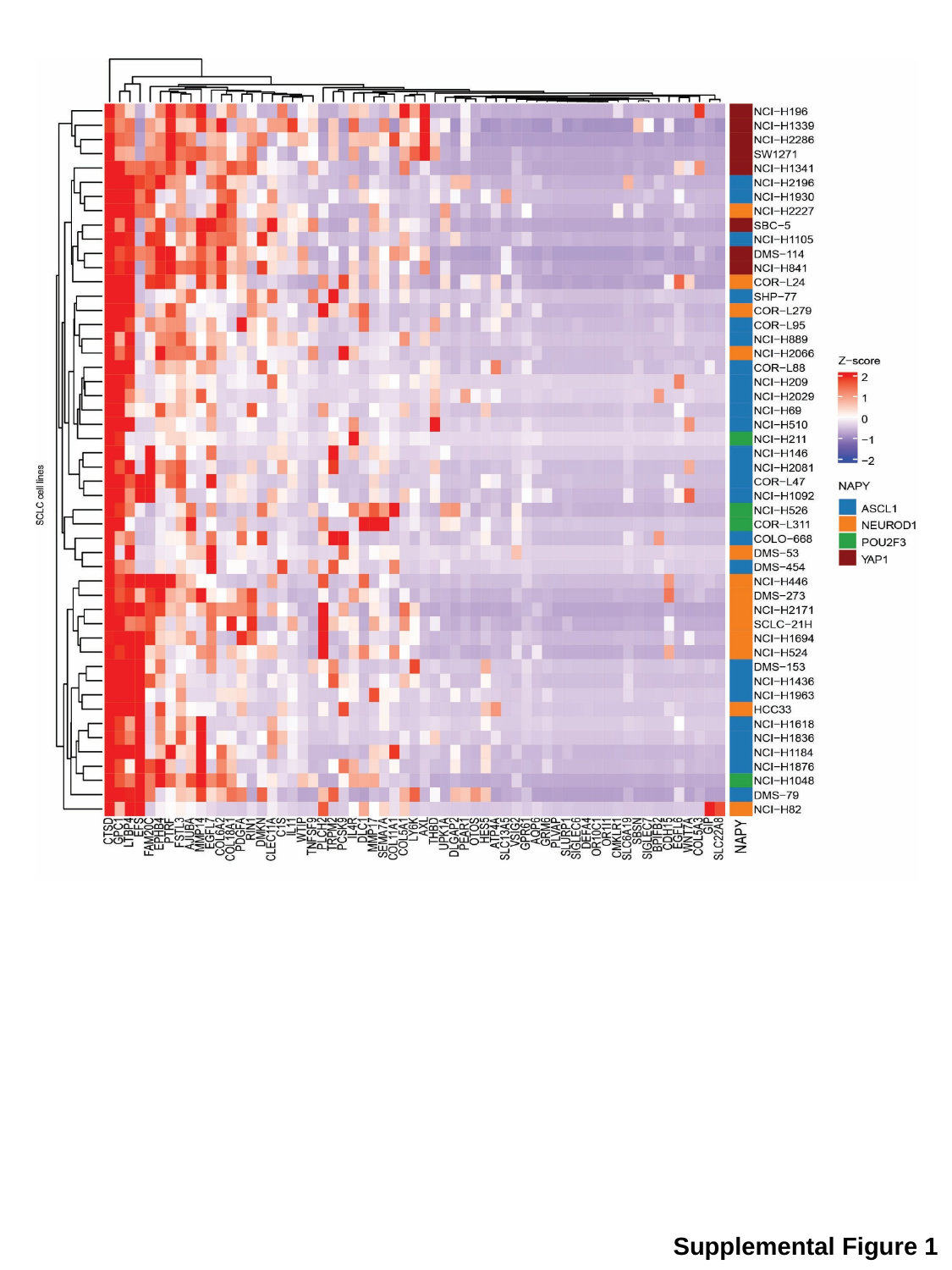

Supplemental Figure 1

### Slide 2
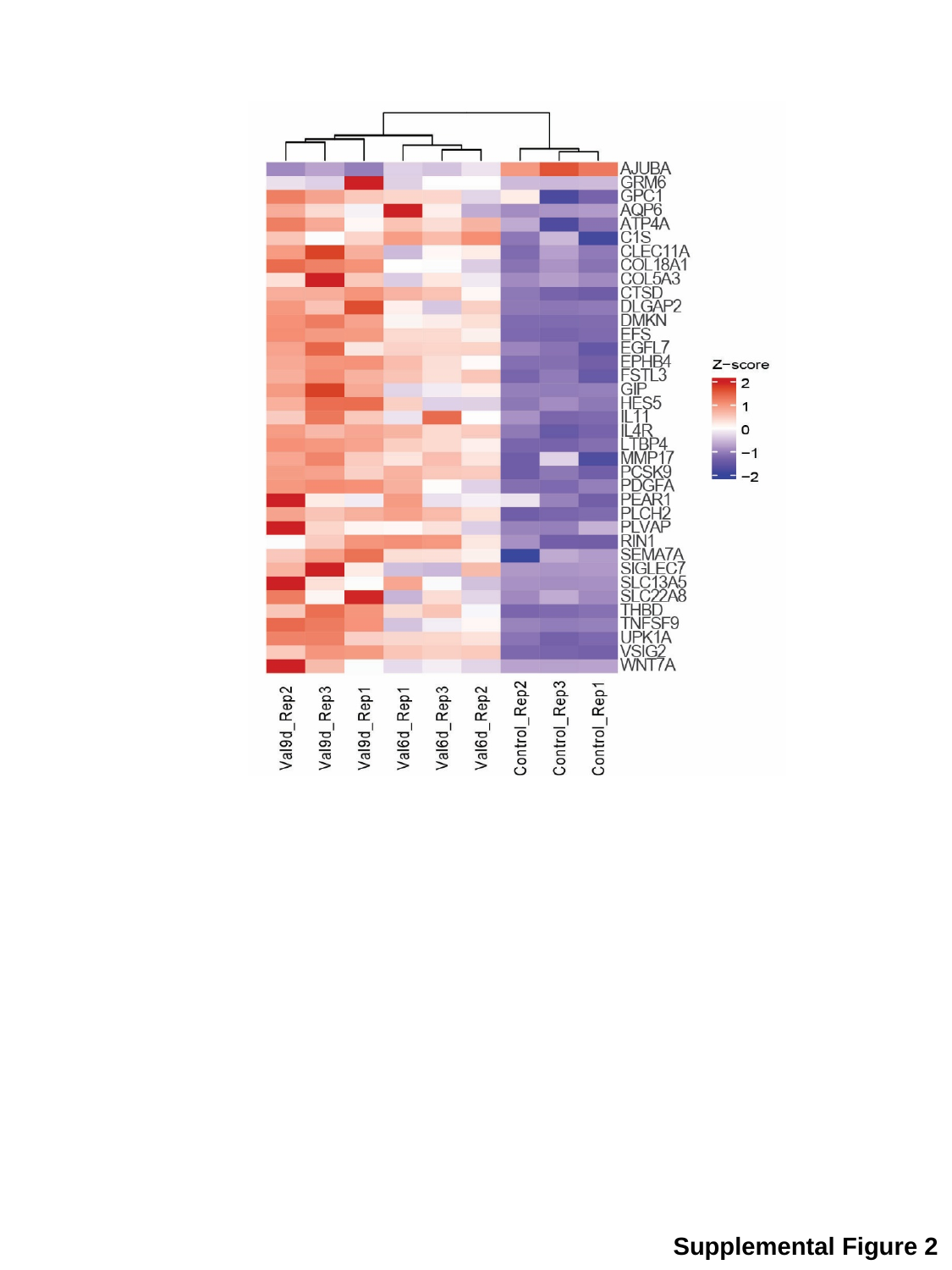

Supplemental Figure 2

### Slide 3
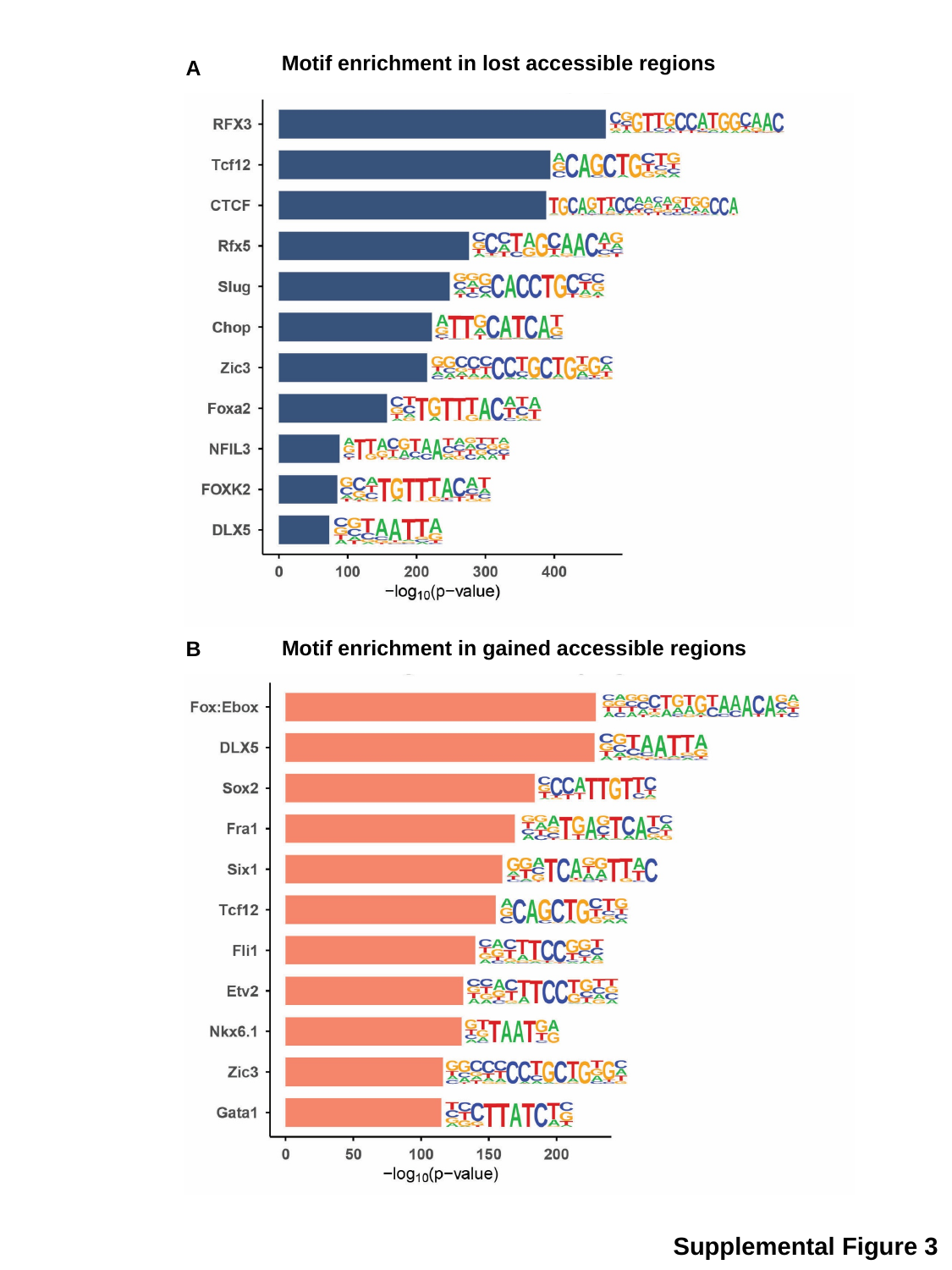

Motif enrichment in lost accessible regions
A
Motif enrichment in gained accessible regions
B
Supplemental Figure 3

### Slide 4
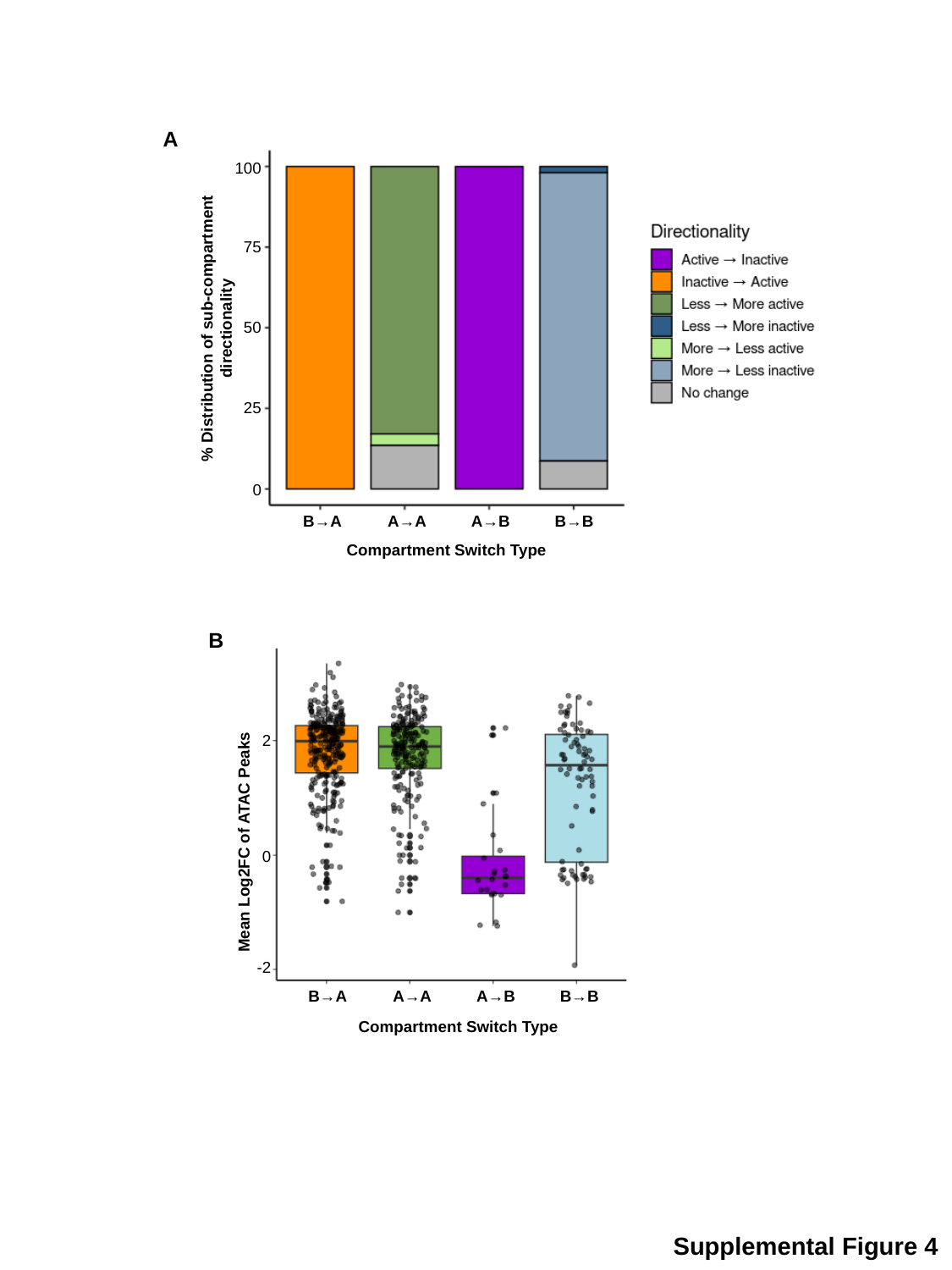

A
100
75
% Distribution of sub-compartment directionality
50
25
0
B→A
A→A
A→B
B→B
Compartment Switch Type
B
2
Mean Log2FC of ATAC Peaks
0
-2
B→A
A→A
A→B
B→B
Compartment Switch Type
Supplemental Figure 4

### Slide 5
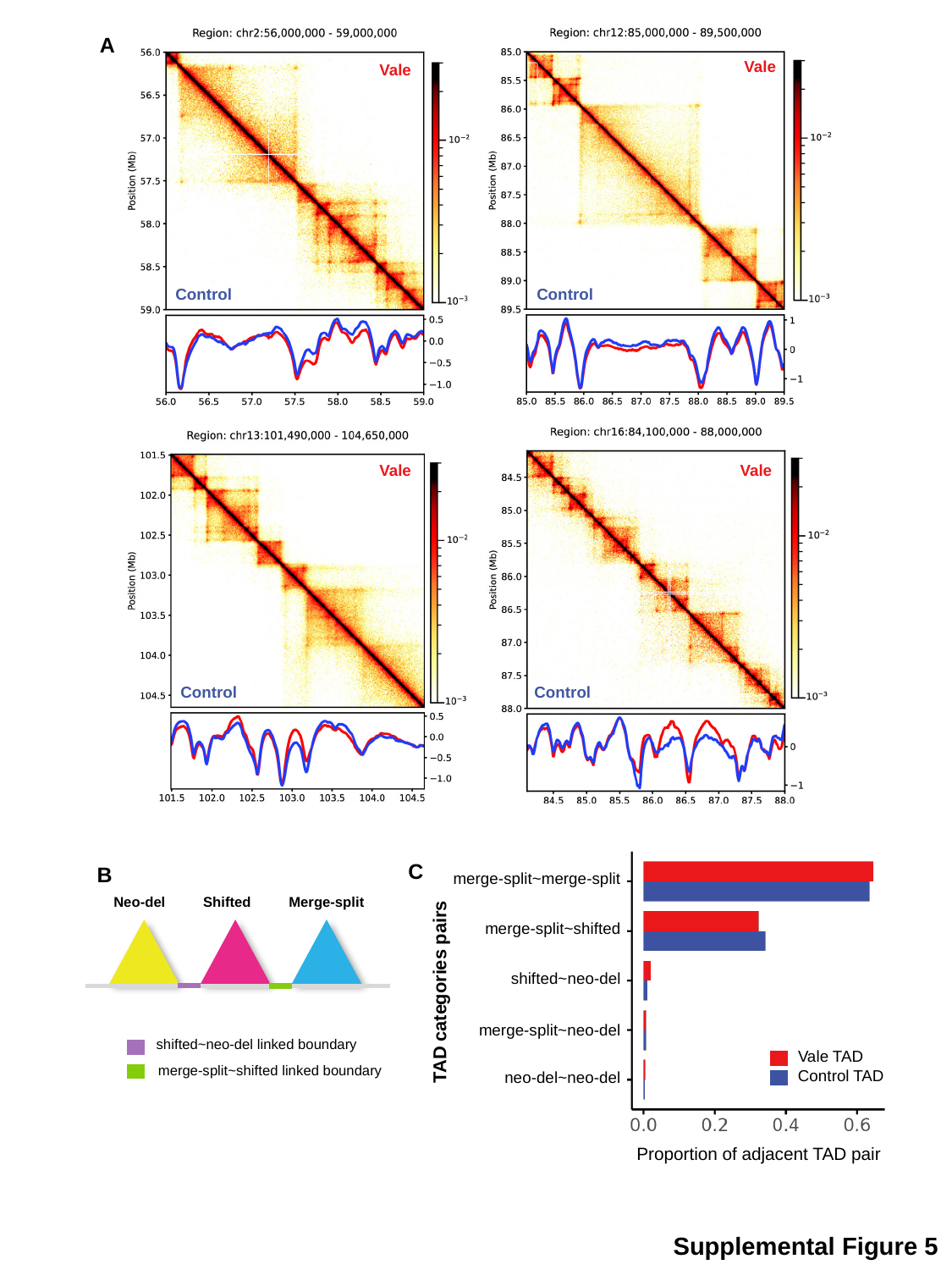

A
Vale
Vale
Control
Control
Vale
Vale
Control
Control
merge-split~merge-split
merge-split~shifted
shifted~neo-del
merge-split~neo-del
neo-del~neo-del
Vale TAD
Control TAD
Proportion of adjacent TAD pair
C
B
Neo-del
Shifted
Merge-split
shifted~neo-del linked boundary
merge-split~shifted linked boundary
TAD categories pairs
Supplemental Figure 5

### Slide 6
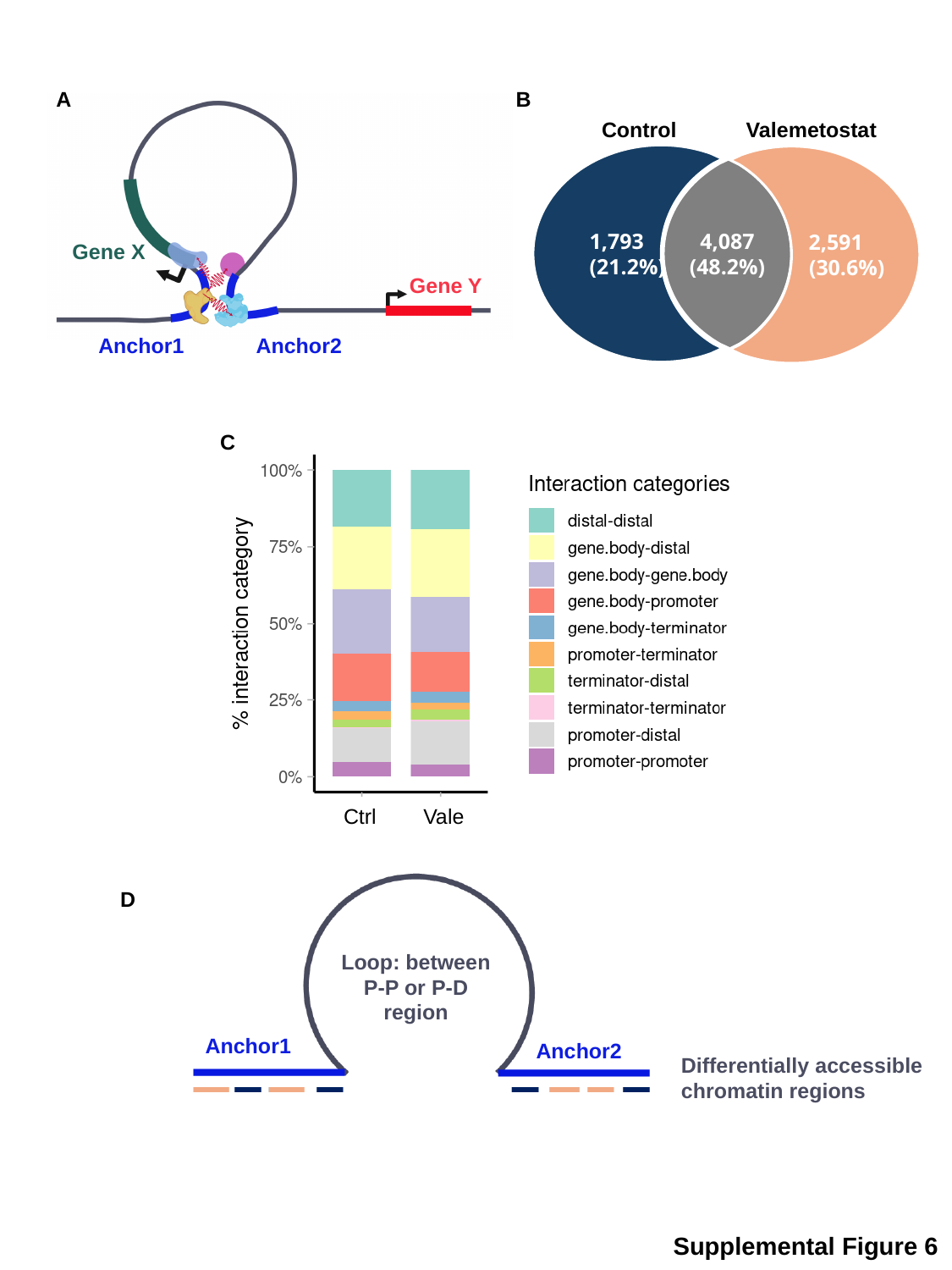

A
B
Gene X
Gene Y
Anchor2
Anchor1
Control
Valemetostat
 2,591
 (30.6%)
4,087
(48.2%)
 1,793
 (21.2%)
C
Vale
Ctrl
Loop: between P-P or P-D region
Anchor1
Anchor2
Differentially accessible chromatin regions
D
Supplemental Figure 6
