## Supplementary material for "EZH2 Inhibition Reshapes 3D Chromatin Architecture to Induce Immunogenic Phenotype in Small Cell Lung Cancer": N/A

**Supplemental Figure Legends:**

**Supplemental Figure 1:** Heatmap depicting the expression patterns of 61 genes, previously reported to be epigenetically silenced by the PRC2 complex in small cell lung cancer (SCLC) cell lines, compared with normal airway epithelial cells across 50 SCLC cell lines from the Cancer Cell Line Encyclopedia (CCLE) database. SCLC molecular subtype annotations (NAPY classification) were obtained from SCLC-CellMiner.

**Supplemental Figure 2:** Heatmap of differentially expressed genes (DEGs) in Valemetostat-treated H146 cells that overlap with the PRC2-silenced gene set reported by Sato *et al*. Of the 61 genes, 32 were differentially expressed at day 6 post-treatment, and an additional 5 genes were identified by day 9 post treatment, resulting in a total of 37 of 61 genes showing differential expression in Valemetostat-treated H146 cells compared with control.

**Supplemental Figure 3:** HOMER *de novo* motif discovery analysis of differentially accessible regions identified by the CoBRA workflow, classified as **(A)** lost or **(B)** gained accessibility based on a cutoff of |log_2_ fold change| ≥ 1 and adjusted p-value < 0.05. Lost regions were defined as peaks with log_2_ fold change ≤ −1, and gained regions as peaks with log_2_ fold change ≥ 1. Each set of regions was independently subjected to motif enrichment analysis using HOMER (findMotifsGenome.pl). The top 11 enriched motifs in each category are shown, ranked in decreasing order of statistical significance (p-value).

**Supplemental Figure 4: (A)** Percentage stacked bar plot showing the distribution of sub-compartment states across differential compartment regions. Sub-compartments are ordered by transcriptional activity according to the following hierarchy A3 > A2 > A1 > A0 > B0 > B1 > B2 > B3, where A3 denotes the most transcriptionally active compartment and B3 denotes the most inactive compartment. Transitions toward increased activity (less → more active) include shifts from lower- to higher-activity A sub-compartments (e.g., A0 to A3), whereas transitions toward increased repression (less → more inactive) include shifts from weaker to stronger B sub-compartments (e.g., B0 to B3). **(B)** Box plots showing the mean differential accessibility scores of ATAC-seq–identified differentially accessible regions that overlap with either matching compartments (A→A and B→B) or switching compartments (A→B and B→A) in day 9 Valemetostat-treated cells. The center line denotes the median, and the box represents the interquartile range (IQR).

**Supplemental Figure 5:** **(A)** Chromatin interaction matrix for selected genomic regions, with the upper diagonal representing the day 9 Valemetostat-treated sample and the lower diagonal representing the control sample. Topologically associating domains (TADs) are indicated as triangles along the diagonal. The bottom panel displays the log2 insulation scores for Valemetostat-treated (red) and control (blue) samples; local minima (dips) indicate TAD boundaries and coincide with TAD borders. **(B)** Schematic depiction of adjacent TADs flanking differential boundary regions, with purple boundaries representing connections between neo-del and shifted TAD pairs, and green boundaries delineating shifted and merge-split TAD pairs. Differential boundary analysis was performed by comparing the log2 insulation score ratios between conditions using the quaich workflow, with a fold-change threshold of 5. We identified 2,202 out of 4,939 total detected boundaries as differential in Valemetostat-treated samples, and 2,148 out of 4,886 total detected boundaries as differential in control samples (relative to Valemetostat-treated samples). **(C)** Grouped bar plot illustrating the proportion of adjacent TAD pair types across differential boundary regions.

**Supplemental Figure 6: (A)** Schematic illustration of promoter-distal (P-D) chromatin loops identified by micro-C. Loops connect promoter regions with distal regulatory elements (often hundreds of kb apart), mediated by DNA-bound transcription factors and co-activators. Interactions are captured through chromatin crosslinking, followed by micrococcal nuclease (MNase) digestion and proximity ligation. **(B)** Venn diagram showing the classification of all identified chromatin loops (within a ≤255 kb width cutoff) into three categories when comparing Valemetostat-treated samples to control: Conserved (present in both, ≈50% of total), Gained (present only in Valemetostat, ≈30% of total), and Lost (present only in control, ≈21% of total). **(C)** Bar plot illustrating the distribution of chromatin loop interactions (for both control and Valemetostat treatments) across various genomic features (e.g., promoter, distal, gene body, terminator). **(D)** Schematic illustration showing the overlap between chromatin loop anchors and differentially accessible peaks identified using the CoBRA workflow. Orange peaks represent regions of gained accessibility (positive log_2_ fold change), and blue peaks represent regions of lost accessibility (negative log_2_ fold change) in day 9 Valemetostat-treated samples compared to untreated controls. Differential accessibility scores were subsequently averaged across anchor pairs (P-P and P-D) for each gene to evaluate the contribution of chromatin accessibility changes to loop formation and gene expression regulation.
